# Modified protocol for quick and large-scale transformation of *Pichia pastoris*

**DOI:** 10.1101/337592

**Authors:** Ravinder Kumar

## Abstract

Over the last couple of decades, methylotrophic yeast, *Pichia pastoris* emerges as an important yeast species owing to its increasing application in industry and basic biological research. Transformation of *Pichia pastoris* cells for the introduction of the gene of interest is common practice for expression and purification of a heterologous protein(s). Presently available protocol of *Pichia pastoris* transformation involves preparation of competent cells and followed by their transformation. Preparation of competent cells requires growth of cells to certain cell density which requires lots of resource, space, time and efforts. This limits the number of transformations that can be performed by an individual at a time. In the present paper, I will describe a modification in the available protocol which makes *P. pastoris* transformation hassle free. The present procedure does not require growth of pre-culture or growth of cells to certain cell density rather cells are grown in a patch on YPD plate(s) and rest procedure is performed in small eppendrof tubes which allow a large number of transformations in quickest possible time with minimal resource and efforts. In the end, I also compare various protocols in tabular form which allows the user to choose best suitable procedure depending on the available resource, time, number of transformations, requirement, and efforts. The present modified protocol does not require big centrifuge and shaker which further makes this procedure more useful. I believe that present protocol of transformation with its many unique features will be really helpful to those working with *P. pastoris.*

## 1 INTRODUCTION

Methylotrophic yeast, *Pichia pastoris* is now an important model in both basic biological research (Gasser et al., 2013) as well as an important industrial yeast species for expression of heterologous proteins (Cereghino et al., 2000; Daly et al., 2005). Apart from that *P. pastoris* is also used in health care industries where this yeast species is used for expression of biopharmaceutical proteins (Daniel et al., 2013). Although budding yeast, *S. cerevisiae* is widely used in basic science research but for many studies, *P. pastoris* remains a choice of model. For example, processes like pexophagy and peroxisome biogenesis are best studied in *P. pastoris* (Farré et al., 2008; Farré et al., 2017). Apart from that *P. pastoris* offers other benefits like the low level of genetic redundancy, can grow on simple media with methanol as a sole carbon source, culture can reach very high density, some of the genes which were absent in budding yeast is present in *Pichia pastoris* (example Atg37) (Nazarko et al., 2014). Apart from that *P. pastoris* processes many features which make it important industrial species. For example, as mentioned above, it can grow to high cell density, can use methanol as a sole carbon source (thereby checking the growth of other unwanted microbial species), glycosylation pattern of proteins is closer to humans (in *S. cerevisiae* proteins are generally hyper glycosylated), availability of many methanol inducive promoters, secretes the low level of endogenous proteins (Cereghino et al., 2000), and more recently availability of more refined complete genome sequence (Sturmberger et al., 2016) of this species.

Efforts were made to introduce DNA of interest for either introduction or deletion of the gene of interest both from perspectives of basic science as well as industrial importance. As a result, several protocols were devised for the introduction of DNA of interest in *P. pastoris* cells including electroporation (Becker & Guarante, 1991), alkali cation method (Ito et al., 1983), treatment of cell with polyethylene glycol (or PEG) (Dohmen et al., 1991) and method involving spheroplast generation method (Cregg et al., 1985). Each of these methods has its own benefits and shortcomings. But all the available procedure for *P. pastoris* transformation involves two steps *viz* 1) preparation of competent cells and 2) transformation of competent cells. Although, electroporation is quick, reliable and allows a large number of transformations, but the preparation of competent cells is a long procedure which requires over-night pre-culture, dilution of pre-culture followed by growth of cells to certain cell density (in all available methods preculture is diluted to cell density ranging from OD_600nm_ 0.2-0.3 followed by growth of cells to OD_600nm_ 0.8-1.2) which may require 4 to 6 hours depending upon strain (Shixuan & Letchworth, 2004). Apart from that, each transformation requires handling of 50-100 mL of media which limits the number of competent cells (different strain) which can be prepared by an individual at a time. Although condensed protocol of *P. pastoris* transformation somehow makes procedure a bit simple (Cereghino et al., 2005) but a modification which can still make the whole procedure of *P. pastoris* transformation more simple, short, quick and reliable is always desired. In all available protocols for *P. pastoris* transformation, preparation of competent cells still remains a limiting and effort intensive step. Therefore, a protocol which does not required competent cell preparation or allows preparation of a large number of competent cells in shortest possible time with a minimum resource, space, time, efforts and does not involve handling of a large volume of culture is always desirable.

In the present study, I am reporting a modification in available protocol for routine transformation of *P. pastoris* which gives a significant number of colonies with a high rate of positive transformants. The present procedure allows transformation without growing cells in tubes and flasks which makes the present protocol simple, quick, reliable allowing transformation of large numbers of samples which is not possible with the presently available procedure for *P. pastoris* transformation. In the end, I also compare present protocol with the already available procedure and discuss merits of the present procedure. The present modified procedure with its unique way of competent cell preparation truly condensed the whole procedure of *P. pastoris* transformation. Further, the present procedure has been tested on different *P. pastoris* background several times over the span of six months. I believe that present modified protocol will be really helpful for those working with *P. pastoris* and required a large number of transformations on a daily or routine basis which is pretty common in industrial settings.

## 2 MATERIAL AND METHODS

### 2.1 Chemical and reagents

DTT (from Roche), yeast extract (Difco), peptone (Difco), dextrose (Fisher), YNB (Bacto), sorbitol (Fisher), restriction enzyme (NEB), electroporation cuvette (Genesee scientific), agarose (Apex), 1 kb Plus DNA ladder (Thermo fisher), 6X DNA loading dye (NEB), pre stained protein marker (Thermo fisher), nitrocellulose membrane (GE), ECL reagents (GE), Tris base (Sigma), CSM-His (Sunrise Science), anti-GFP antibodies (Clonetech, JL-8), goat anti HRP conjugated secondary antibodies (Biorad), SDS (Fisher), NaOH (Fisher), glycerol (Calbiochem), TCA (Sigma), methanol (Fisher), electroporation unit (BTX, ECM20, version 1.04) bench top centrifuge (Eppendrof), antibiotics (sigma), UV-visible spectrophotometer (from Beckmann Coulter), nonfat skimmed milk powder (Apex), X-ray film processor (from Knoica SRX-101A), autoradiography films or sheets (Blue Devil). All other reagents were of either analytical or molecular grade.

### 2.2 Strain and plasmids

All the *Pichia pastoris* strains used in the present study are isogenic to the PPY12h background. Cartoon presentation of plasmid (showing essential elements) integrated into *P. pastoris* genome for introducing gene(s) of interest using the protocol described in present manuscript is shown in figure 2.1.

### 2.3 Media

YPD media (2 % yeast extract, 1 % peptone and 2 % dextrose). SD+CSM-His (0.17 % YNB without amino acid and ammonium sulfate, 0.5 % ammonium sulfate, 0.08 % CSM-His, 2 % agar, 2 % dextrose), YPD + Zeocin (YPD + 100 μg/mL Zeocin). All cultures were grown at 30 °C, 250 rpm.

### 2.4 Protein extraction

Transformants were patched on fresh selection plate(s) and plates were incubated at 30 °C till patches grow (generally 1-2 days). One mL YPD media were inoculated using cells from the patch. Tubes were incubated at 30 °C, 250 rpm. Next day OD_600nm_ of cell suspension was checked and cell suspension equal to one OD was transferred into fresh 1.5 mL eppendrof tube and TCA (Trichloroacetic acid) was added to the tube such that final concentration of TCA was around 12.5 %. Tubes were incubated at -80 °C for one hour. After one-hour tubes were taken out from deep freezer, thawed at room temperature and vortexed for 30 second. The tube was then centrifuged at 18000 g for 8 min and the supernatant was discarded. The resulting pellet was resuspended in 1 mL of chilled 100 % acetone (using water bath). Tube(s) were again centrifuged as above, and the supernatant was discarded carefully without disturbing or losing protein pellet. The protein pellet was air dried and resuspended in 100 μL Laemmli buffer (Laemmli, 1970).

### 2.5 Immunoblotting

Whole cell lysates (extracted above) were resolved on 10 % SDS-PAGE and proteins were transferred onto the nitrocellulose membrane as described elsewhere (Towbin et al., 1979). Efficiency and quality of transfer were checked by staining the blots with Ponceau S stain just before incubating the blots in blocking solution (5 % nonfat skimmed milk powder in TBST) (Romero-Calvo et al., 2010). Proteins were detected by using monoclonal anti-GFP and HRP conjugated goat anti-mouse as secondary antibodies respectively. Blots were developed using chemiluminescence (from GE).

### 2.6 Fluorescence microscopy

Images were captured and analyzed using a fluorescence microscope and software as described elsewhere (Wang et al., 2017).

### 2.7 Transformation protocol

The detailed protocol for *Pichia pastoris* transformation is described below. On a fresh YPD plate, a patch of required strain(s) was prepared, and plate(s) were incubated at 30 °C. Patch size of around 2 cm by 1.5 cm (length × breadth) was sufficient for two transformations (giving around 15-18 OD_600nm_ cells). After 18-24 hour of incubation, cells from the patch(s) can be used for transformation. Transfer 1 mL of YPD in 1.5 mL sterile Eppendrof tube. Add 40 μL of DTT (from a stock of 1 M, prepared from DDT powder from Roche) and 40 μL HEPES-NaOH buffer (from a stock of 1 M pH8). Scrap the cells from the patch with help of 200 μl sterile tips or blunt ended sterile toothpicks and transfer them in a tube having YPD with DTT and HEPES buffer. Make sure that cells are resuspended completely. Shake the tube gently for 15 min at 30 °C. After 15 min of gentle shaking at 30 °C, wash the cells twice with sterile water. During washing steps, cells should be pelleted down at 3000 g for 3 min. After completing washing steps, incubate the tube on ice for 3-5 min, mix with DNA (PCR product or digested plasmid) and gently mix the content of tube. Transfer the content of tube in pre-labeled electroporation cuvettes which were already kept on ice. After transfer content in cuvette give the electric pulse to the cells at the following settings (Voltage: 1500 VH, Resistance: 200 Ω, Capacitor: 0025 μF using BTX, ECM20, version 1.04). Just immediately after the electric pulse, add 1 mL ice-cold 1 M sorbitol (as recovery medium) into the electroporation cuvette and mix well. If the selection is on an antibiotic plate, incubate the cuvette with cells at 30 °C for 2-3 hour and if a selection is on dropout plate, plate the content of tube just after the addition of recovery medium. Note that steps which can be modified in present modified procedure are discussed in the discussion section.

## 3 RESULTS

### 3.1 Need for a new or modified protocol

As mentioned in the introduction that, all the available protocol for *P. pastoris* transformation involved over-night pre-culture (source of inoculum), dilution of pre-culture followed by growth of cells to OD_600nm_ close to 1.2 which take around 5-6 hour depending upon strain(s) (table 1 showing comparison of various available protocol) (Ito et al., 1983; Dohmen et al., 1991; Cregg et al., 1995; Cereghino et al., 2005; Shixuan & Letchworth, 2004; Hinnen et al., 1978). Further, steps involved in preparing competent cells again require 1-2 hour depending upon the number of strains handled at a time. And if the selection is on an antibiotic plate, it again increases the time till final plating of cells on plate. This means whole day is required to complete the transformation experiment. Apart from that present protocols involve growing cells in 50-100 mL media which again limit the number of flasks which can be handled by an individual at a time. In short transformation of *P. pastoris* is a lengthy process requiring a lot of efforts and resource. Basic steps of different protocols for *Pichia pastoris* transformation are compared in table 1.

**Table 1.** Comparison of present modified protocol with previously published protocols form economic point of view.

| **<u>Heat Shock</u>**<br>(Dohmen et al., 1991)<br>[A] | **<u>Alkali Cation</u>**<br>(Ito et al., 1983)<br>[B] | **<u>PEG</u>**<br>(Klebe et al., 1983)<br>[C] | **<u>Electroporation</u>**<br>(Becker & Guarante, 1991)<br>[D] | **<u>Condensed Protocol</u>**<br>(Cereghino et al., 2005)<br>[E] | **<u>Present study (modified electroporation)</u>** |
| --- | --- | --- | --- | --- | --- |
| Growth of preculture | Growth of preculture | Growth of preculture | Growth of preculture | Growth of preculture | Making of patch on plate |
| Dilution of preculture | Dilution of preculture | Dilution of preculture | Dilution of preculture | Dilution of preculture | Scrap the cells from patch and resuspend them in 1 mL YPD with DTT and HEPES pH8 |
| Growth of cells to desired OD<sub>600nm</sub> (0.6 to 1.0) | Growth of cells to desired OD<sub>600nm</sub> | Growth of cells to desired OD<sub>600nm</sub> | Growth of cells to desired OD<sub>600nm</sub> | Growth of cell to desired OD<sub>600nm</sub> | Incubate the cells at 30 °C, for 15 min in gentle shaking condition |
| Harvesting culture and washing with solution/buffer | Harvesting culture and washing with solution/buffer | Harvesting culture and washing with solution/buffer | Harvesting culture and washing with solution/buffer | Harvesting culture and washing with solution/buffer | Wash the cells twice with sterile 1 mL water each time. |
| Resuspend cells in 0.02 volume of above same solution | Incubate the cells with buffer for 1 h at 30°C. | Resuspend cells in buffer | Suspend the cells in 100 mL of YPD medium plus HEPES, add 2.5 mL of 1 M DTT and gently mix | Resuspending of cells in 9 mL BEDS solution supplemented with 1 mL 1.0 M DTT | Discard supernatant |
| Freeze cells or proceed for transformation | Mix vector DNA with cells | Add DMSO and freeze or proceed for transformation | Incubate at 30°C for 15 min | Incubate the cells for 5 min, 100 rpm, 30 °C | Mix DNA with cells |
| Mix DNA with cells | Mix carrier DNA with cells | Mix DNA with cells | Bring to 500 mL with cold water | Incubate the cells for 5 min, 100 rpm, 30 °C | Incubate cells on ice for 3-5 min |
| Add 40% polyethylene glycol (PEG), | Incubate samples at 30°C for 30 min | Mix carrier DNA with cells | Again, wash the cells | Harvest the cells at 500 g, 5 min, RT and resuspend them in 1 mL BEDS solution without DTT | Give electric pulse to cells |
| Incubate for 60 min at 30°C | Add 0.7 mL of PEG + LiCl solution, and briefly vortex to mix | Incubate samples in a 37°C water bath for 5 min. | Freeze at -70 °C or proceed for transformation | Freeze at -70 °C or proceed for transformation | Either plate the cells or incubate depending upon selection |
| Heat shock at 42°C for 10 min | Incubate samples at 30°C for 30 min | Add another solution and mix well | Mix DNA with cells | Mix DNA with cells | Mix the cells with recovery medium |
| Pellet cells | Heat-shock at 37°C for 5 min | Incubate tubes in a 30°C water bath for 1 h | Incubate on ice | Incubate on ice | Incubate for 2-3 hour or spread on desired plate |
| Resuspend cells in buffer | Centrifuge samples at 2000g and resuspend in 0.1 mL of H<sub>2</sub>O | Pellet down cell and resuspend in another buffer (repeat this again) | Giving electric pulse to cells | Giving electric pulse to cells |  |
| Repeat last two steps | Centrifuge samples at 2000g and resuspend in 0.1 mL of H<sub>2</sub>O | Again, pellet down cell and resuspend in another buffer | Mix the cells with recovery medium | Mix the cells with recovery medium |  |
| Incubate for 2-3 hour or spread on desired plate | Incubate for 2-3 hour or spread on desired plate | Incubate for 2-3 hour or spread on desired plate | Incubate for 2-3 hour or spread on desired plate | Incubate for 2-3 hour or spread on desired plate |  |

Note: In this table only basic steps are compared without describing fine details of each step(s), buffer used and their composition. For this readers are advised to check original proper (reference provided).

Therefore, a protocol which is short, reliable and robust which can reduce the efforts, time, resource and allows transformation of a large number of strains will be important. Importantly, a protocol which does not require growth of cells in flasks or handling of bulk culture i.e. skip culturing step is always desirable. Therefore, in the present study, I will be describing a protocol for *P. pastoris* transformation which does not require culturing the cells and allows a large number of transformations with minimum efforts and also requires less resource in terms of use of lab equipment like shaker, big centrifuge, and other lab reagents.

### 3.2 Basic workflow

A detailed procedure of *Pichia pastoris* transformation is described in materials and methods section. Here only the basic workflow of the protocol is described, and comparison was drawn between the protocol described in the present study and other available protocols (Figure 1). Basic elements of plasmid integrated in *P. pastoris* using present modified protocol is shown in figure 2A. The present protocol does not require growth of pre-culture and growth of cells in big flasks. A patch of around 2 cm by 1.5 cm is sufficient for two transformations (Fig. 2B). This saves lots of media used in pre-culture and growing cells. This step also reduces use of plastic wares (culture tubes) and proved to be more economical. Required strain(s) are patched properly on fresh YPD plate and plate(s) were incubated at 30 °C for 18-24 hour. This step allows transformation even in absence or non-availability of big shakers. The appearance of a fine layer of cells in the patched area suggested that patch is ready for transformation. Transfer 1 mL of YPD with 40 mM DTT, 40 mM HEPES buffer pH8 in a required number of sterile eppendrof tubes. Using a sterile 200 mL tips or toothpick scrap the cells and resuspended in YPD. Make sure cells are dispersed properly. One can vortex the tube to disperse cells to get uniform cell suspension. Gently shake the tube for 15 min at 30 °C. After completion of this step, cells were washed twice with sterile water. Each time cells were pelleted down by centrifugation at 3000 *g* for 3 min. This allows preparation of competent cell more quick, easy and cost effective as it does not require big tubes (50 mL tubes) to harvest cells, washing of cells in different buffers. The supernatant was discarded, and the cell pellet was resuspended in such a way that total volume of cell suspension was around 50-70 μL and a required DNA (intact plasmid, digested plasmid or PCR product) was mixed properly with cell suspension and the cell suspension was transferred into an electroporation cuvette. Electric pulse or shock was given using settings described in material and methods. After electric shock 1 mL recovery medium (1 M sorbitol or 2 % glucose) was added to cuvette and cells were mixed well. The content of cuvette was transferred into eppendrof tube and cells were pellet down by centrifugation at 3000 *g* for 3 min. The supernatant was discarded, and the cell pellet was resuspended in sterile water such that final volume is no more than 100 μL. Cells were plated on a required plate.

**Figure 1.**
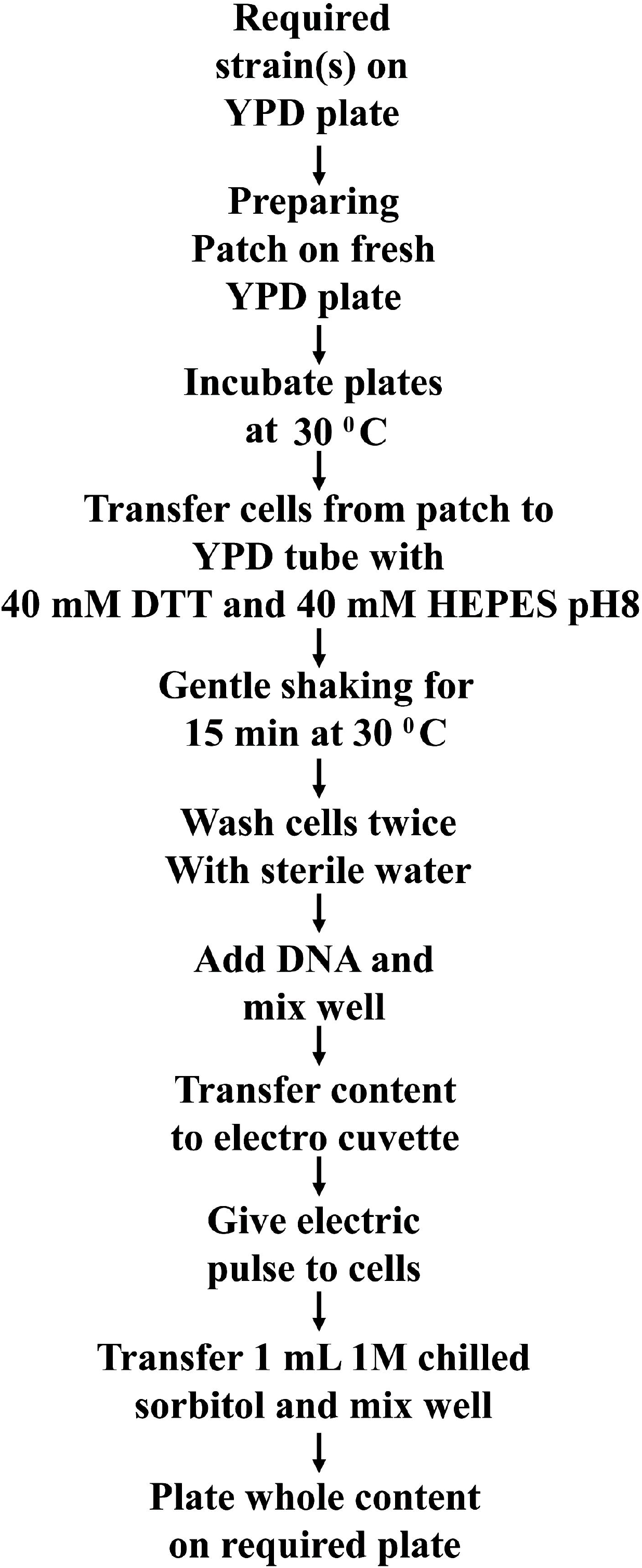
Schematic showing basic workflow of present modified protocol for Pichia pastoris transformation.

**Figure 2.**
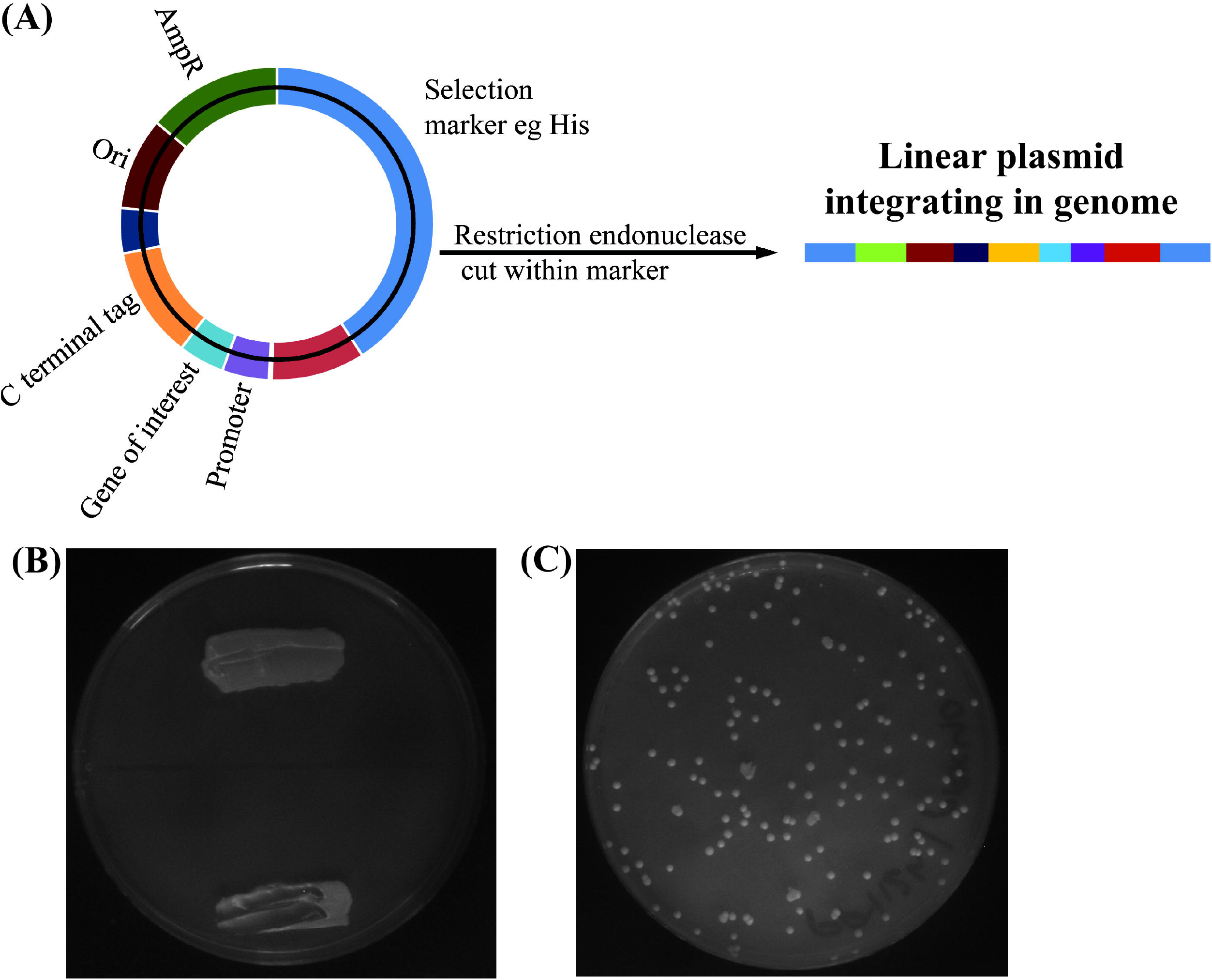
Transformation of *P. pastoris* using modified protocol. (A) Basic features of plasmid integrated in *P. pastoris* gene using present procedure. (B) YPD plate showing size of patch used for transformation. (C) Image of one of the transformant plate. Note number of transformants (colonies) on transformant plate vary 3-5 times compared to one shown in figure 2C.

Number of transformants that appeared on transformant plate vary significantly and represented image of one of the transformant plate is shown in figure 2C. Cost effectiveness of present modified procedure for *P. pastoris* transformation is shown through table 2.

**Table 2.** Comparison of various published protocol for *P. pastoris* transformation with present modified procedure

| <u>**Present procedure**</u> | <u>**Other published protocol**</u><br>{A to E}* | <u>**Comments**</u> |
| --- | --- | --- |
| No requirement of culture tube for pre-culture | Require culture tube for preculture | Present procedure is cost effective |
| 45-50 mL YPD agar is sufficient for 10 transformations | Relatively more media is required | Present procedure is cost effective |
| Require incubator | Require big shaker, more electric power | Present procedure is cost effective |
| Cell preparation is performed in 1.5 mL tube | May require bigger tubes (50 mL, 10 tubes) | Present procedure is cost effective |
| Small benchtop centrifuge is sufficient | Since high volume of media and buffer handling is required which necessitates big centrifuge { | Present procedure is cost effective |
| No special buffer for cell washing as washing is done with water | Require special buffer depending upon procedure, may require different buffers at different washing steps | Present procedure is cost effective |
| Less requirement of power for different steps | More power is required at different steps | Present procedure is cost effective |
| Relatively less investment in terms of big shaker and centrifuge | Big investment in terms of big shaker and centrifuge | Present procedure is cost effective |

\***Note:** In this table comparison is made based on requirement on different steps in previously published protocols (A to E in table 1).

### 3.3 Integration and expression of a gene of interest

Working of the protocol was checked by introducing PpCUE5-GFP and PpPGK1-GFP. Pgk1 is a cytosolic enzyme involved in glycolysis and gluconeogenesis (Hitzeman et al., 1980; Blake and Rice, 1981) while Cue5 is a cytosolic ubiquitin binding protein (Shih et al., 2003; Lu et al., 2014). Both fusion proteins were checked by detecting C-terminal GFP tags using anti-GFP antibodies (Fig. 3A). Western blot image also shows that the expression of PpCue5-GFP and PpPgk1-GFP was similar in all the transformants checked by western blot. Protein loading was shown by Ponceau S stained blot image which clearly showed the amount of protein loaded in each well were similar (Fig. 3B). Out of six colonies checked by western blot, five were positive and was negative for both PpPgk1-GFP and PpCue5-GFP. In both the cases colonies were selected randomly for verification by western blot. Plasmids (with PpCUE5 and PpPGK1 under their native promoter and C-terminal GFP tag) was integrated at *his* locus (Fig. 2A) after linearizing plasmid by *Eco*NI. Introduction and expression of a gene introduced using the present modified procedure of *P. pastoris* transformation were also checked and confirmed by detecting GFP in cells using fluorescence microscopy (Fig. 3C and 3D for PpPgk1-GFP and PpCue5-GFP respectively). Taken together with data of western blot and microscopy data showed that gene(s) introduced by a modified procedure for *P. pastoris* transformation were integrated properly into the genome at required locus or position and able to express properly.

**Figure 3.**
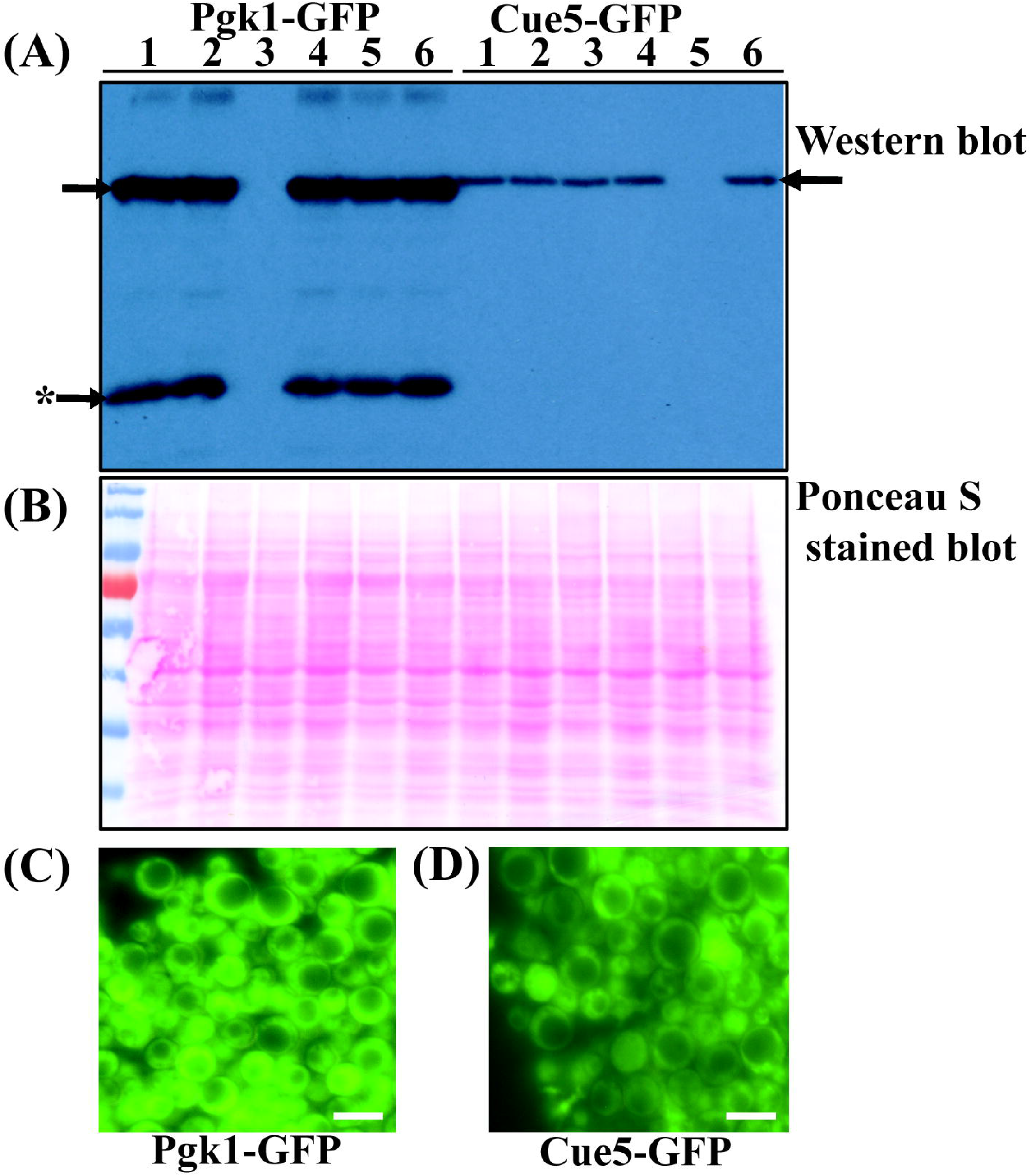
Integration of gene of interest in *P. pastoris* genome. (A) Western blot showing positive transformants for PpPgk1-GFP (first 1-6 lane, left side) and PpCue5GFP (last 1-6 lane, right side) Expected bands are pointed by arrow towards them. Fragmented or nonspecific bands. Mass of both fusion proteins are mentioned in main text. (B) Protein loading is shown by Ponceau S stained blot. Both the fusion proteins were detected at expected size range. Western blot results were also confirmed by detecting GFP signal (C) PpPgk1-GFP and (D) PpCue5-GFP. Scale bar represent 5 μM.

Both western blot as well as microscopy data showed significant difference in abundance of Pgk1 and Cue5 and our present data is accordance with previous studies (Kulak et al., 2014) suggesting that observed difference in protein abundance is not due to experimental artifact. Note in both the case gene is under its endogenous promoter. Mass of fusion protein was 65.7 kDa and 71.6 kDa for Cue5-GFP and Pgk1-GFP respectively.

### 3.4 Factors affecting transformation efficiency

The efficiency of transformation depended on several factors including the physiological state or age of cells or culture, way the competent cells are prepared, nature of recovery medium, the number of cells or cell density, amount of DNA. It is important to study how different factors(s) affect the efficiency of transformation. Therefore, in the present study, I check the effect of age of patch on YPD plate, use of DDT, HEPES buffer during competent cell preparation and recovery medium after giving an electric shock to cells. Presently available data showed that age of patch on YPD plate affect transformation efficiency significantly (Fig. 4A). After 3 days I could get only a few colonies and after five days I could not get any colony on transformants plate (data not shown). Further, it was observed that application of DTT and HEPES pH8 at a final concentration of 40 mM improve the efficiency of transformation significantly (Fig. 4B, C respectively). Just like previous reports DTT and HEPES at 40 mM concentration gives best results, DTT and HEPES more than 40 mM does not increase transformation efficiency significantly. Application of chilled 1 M sorbitol or 2 % YPD as recovery medium does not affect transformation efficiency significantly (fig. 4D). Thus, some factors hardly have any effect on transformation efficiency while others are crucial. Factors like cell density, amount of DNA added to competent cells were not investigated in present study as these factors were already investigated by other lab (Shixuan & Letchworth, 2004) and I believe these may behave similarly in present study. Only those factors were investigated which were unique to present procedure like age of patch on YPD plate. Although it was observed that for high efficiency transformation plasmid should be linearized by digestion from middle of marker used during transformation.

**Figure 4.**
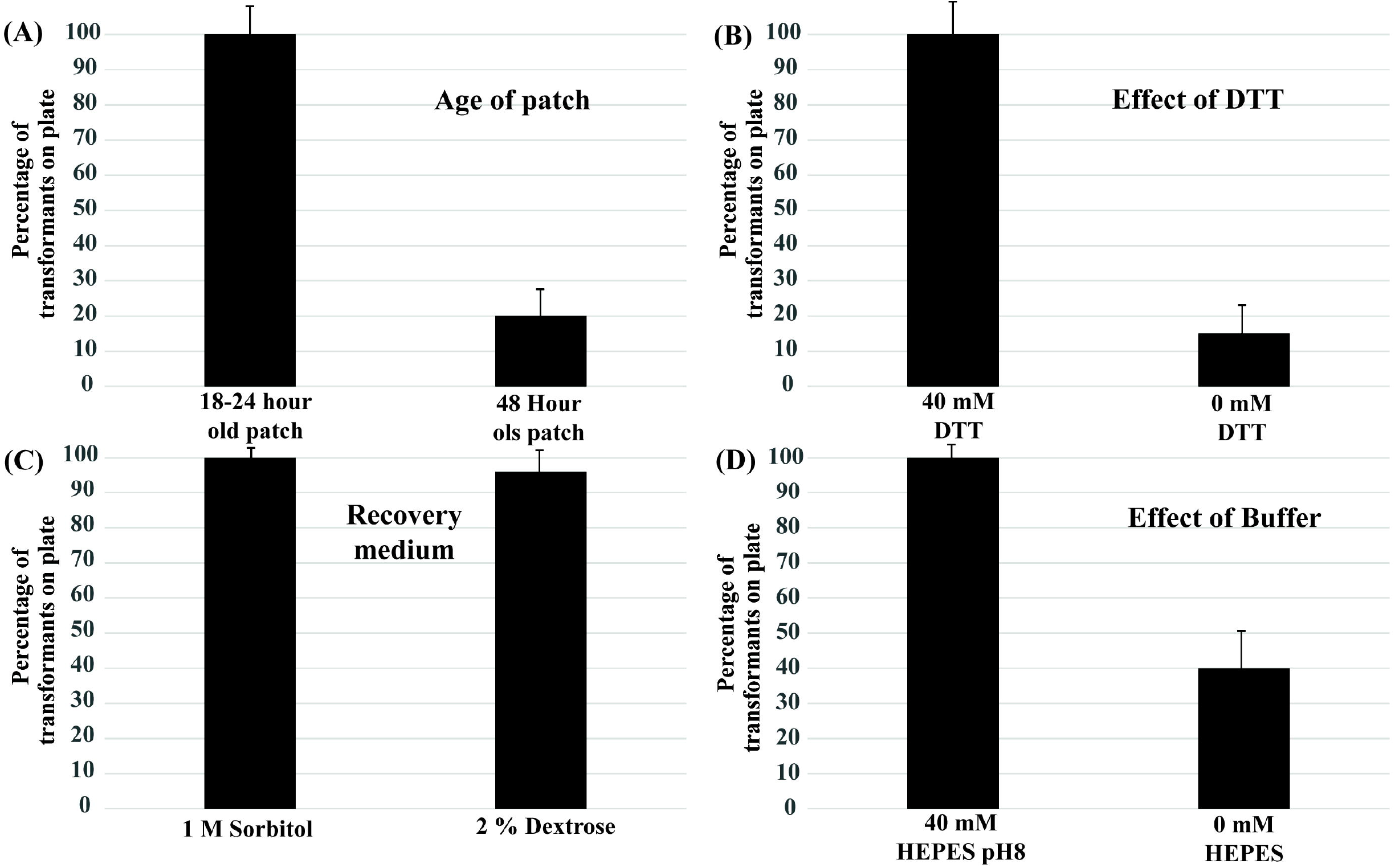
Factors affecting transformation efficiency. Effect of (A) age of patch on YPD plate, (B) effect of reducing agent (DTT), (C) recovery medium and (D) HEPES buffer in on transformation efficiency. Note these parameters were checked on different background *P. pastoris* although data is shown only for PPY12h background.

## 4 DISCUSSIONS

Based on the results and comparison of already published protocol for *P. pastoris* transformation with the procedure described in this paper, it can be said that present procedure got a clear advantage of being short, simple, economical and well suited for large-scale transformation. By getting rid of steps which include overnight pre-culture, dilution of pre-culture and then further growth of cells to certain cell density, the present procedure allows a large number of transformation with minimum efforts, resource and time. Since patches were made on plates which were incubated at 30 °C and the rest steps were carried out in small 1.5 mL eppendrof tubes means the present protocol does not require big shaker and centrifuge which is itself a big advantage especially early days of lab establishment. By simplifying the step of competent cells preparation, the present procedure makes sure that a large number of transformation can be carried out in quickest possible time and added much needed high throughput element in *P. pastoris* transformation which is very important especially in industries where a large number of transformations are carried out on a daily basis. The procedure described in the present paper is also very economical. One patch of 2 cm × 1.5 cm (length × width) is enough for two transformations and one can prepare at least 6 patches of this size means 10-12 transformation can be performed using a single plate with 23-25 mL media compare other available procedure which requires growing of cells in 50-100 mL media for each transformant. Apart from saving on media per transformation, there is a huge saving on other plastic ware, buffers if a large number of transformations are required. Technical improvements in *P. pastoris* transformation are already described by Cai et al. (2001) certainly make *P. pastoris* transformation more robust. I believe that present modified protocol for expression of gene of interest is simpler compared to commercial kits (https://tools.thermofisher.com/content/sfs/manuals/easys_elect_man.pdf) and this may enhance the utility of present procedure.

It is important to mention that efficiency of transformation is affected by nature of plates. It is highly recommended to avoid old plates and does not allow patch grows too thick. It was observed that transformation efficiency falls dramatically as the thickness of patch increases or patch become older. I find that patch generally of 18-24 hour old is most suitable for preparing cells for transformation. It is also advised that one should prepare patch using healthy growing cells. Patch prepared for old dying cells significantly reduces transformation efficiency. Apart from that described procedure will also of great importance for those who do not have access to large shakers and centrifuges. It is important to mention that addition of reducing agent (example DDT) dramatically affects transformation efficiency and present observation is in accordance with the previously published protocol (Cereghino et al., 2005). It was also observed that use of 1 M sorbitol as a recovery medium is not essential and one can also use 2 % dextrose if selection will be made on dropout media lacking required amino acid. But if selection will be made of antibiotic plate, even YPD is good enough as a recovery medium. It is important to mention that number of colonies on transformant plates is highly dependent upon nature of DNA (plasmid or PCR product), gene locus for integration of DNA, nature of selection, number of homologous residues in DNA, whether the transformation is for gene deletion or introduction of gene and so on. It is important to mention that apart from introduction of gene of interest present modified protocol was also suitable for deletion of endogenous gene or ORF (dada not shown).

The present protocol has been tested on different *P. pastoris* background including GS200, GS115, PPY12h and PPY12m over a period of six months. Apart from introducing gene through integrating plasmid at gene locus (generally selection marker like *his, arg),* I was able to integrate the cassette within ORF (of the gene of interest) for N-terminal tagging of protein. Incubation of cells in water with DTT and HEPES pH8 at a final concentration of 40 mM in place of YPD is equally good. Further, in present procedure all steps can be carried out at room temperature without requiring refrigerated centrifuge or ice and incubation of cuvettes after mixing DNA with cells does not affect transformation in a significant way and same goes with the addition of 1 M sorbitol as recovery medium. Although I have not checked the effect of freezing competent cells prepared by present modified procedure, I believe that present procedure makes freezing of competent cells irrelevant and save space in the deep freezer. In the
end, it can be said that present procedure makes the transformation of *P. pastoris* simpler, quick, economical, and less exhaustive and may become choice of method both in labs and industries.

## ACKNOWLEDGMENTS

I regret not being able to refer to the work of everyone in the field. I am thankful to UCSD for providing me space and other necessary facilities which helped me in completing this manuscript. I am equally thankful to Dr. Piyush Kumar from UNT, Texas, for his effort in checking manuscript and valuable and constructive suggestions.

## ORCID

Dr Ravinder Kumar, PhD, https://orcid.org/0000-0003-1922-0030

